# When rarity has costs: coexistence under positive frequency-dependence and environmental stochasticity

**DOI:** 10.1101/161919

**Authors:** Sebastian J. Schreiber, Masato Yamamichi, Sharon Y. Strauss

**Affiliations:** Department of Evolution and Ecology and Center for Population Biology, University of California, Davis; Hakubi Center for Advanced Research, Kyoto University, Sakyo, Kyoto 606-8501, Japan; Center for Ecological Research, Kyoto University, Otsu, Shiga 520-2113, Japan

**Keywords:** positive frequency-dependence, coexistence theory, environmental stochasticity, Allee effects, storage effect, competition, alternative stable states, invasion success, competitive exclusion, niche overlap, reproductive interference

## Abstract

Stable coexistence relies on negative frequency-dependence, in which rarer species invading a patch benefit from a lack of conspecific competition experienced by residents. In nature, however, rarity can have costs, resulting in positive frequency-dependence (PFD) particularly when species are rare. Many processes can cause positive frequency-dependence, including a lack of mates, mutualist interactions, and reproductive interference from heterospecifics. When species become rare in the community, positive frequency-dependence creates vulnerability to extinction, if frequencies drop below certain thresholds. For example, environmental fluctuations can drive species to low frequencies where they are then vulnerable to PFD. Here, we analyze deterministic and stochastic mathematical models of two species interacting through both PFD and resource competition in a Chessonian framework. Reproductive success of individuals in these models is reduced by a product of two terms: the reduction in fecundity due to PFD, and the reduction in fecundity due to competition. Consistent with classical coexistence theory, the effect of competition on individual reproductive success exhibits negative frequency-dependence when individuals experience greater intraspecific competition than interspecific competition i.e., niche overlap is less than one. In the absence of environmental fluctuations, our analysis reveals that (1) a synergistic effect of PFD and niche overlap that hastens exclusion, (2) trade-offs between susceptibility to PFD and maximal fecundity can mediate coexistence, and (3) coexistence, when it occurs, requires that neither species is initially rare. Analysis of the stochastic model highlights that environmental fluctuations, unless perfectly correlated, coupled with PFD ultimately drive one species extinct. Over any given time frame, this extinction risk decreases with the correlation of the demographic responses of the two species to the environmental fluctuations, and increases with the temporal autocorrelation of these fluctuations. For species with overlapping generations, these trends in extinction risk persist despite the strength of the storage effect decreasing with correlated demographic responses and increasing with temporal autocorrelations. These results highlight how the presence of PFD may alter the outcomes predicted by modern coexistence mechanisms.

## Introduction

Understanding mechanisms of multispecies coexistence is one of the central topics in community ecology. Stabilizing forces of niche differentiation (intraspecific suppression being stronger than interspecific suppression) and fitness differences among species are thought to lie at the heart of stable coexistence by species [Chesson, 2000]. When stabilizing forces are sufficiently strong relative to fitness differences, the per-capita growth rate functions of competing species exhibit negative frequency-dependence (NFD) in which the rare species gains a growth rate advantage. These properties have been considered primarily in light of resource competition between species, with fitness functions that give fitness advantages to the rarer species through competitive release. The rarer species escapes intense conspecific competition, while the more common species strongly suppresses itself.

Rare species may, however, experience costs that outweigh the fitness gains of competitive release through a variety of mechanisms involving positive density-or frequency-dependence. Costs due to positive density-or frequency-dependence (PDD or PFD, respectively) include a range of Allee effects (reviewed in [Taylor and Hastings, 2005, Berec et al., 2007]) including reproduction (e.g., a lack of mates when rare [Courchamp et al., 1999, Schreiber, 2003, Zhou and Zhang, 2006]), survival (e.g., a loss of required mutualists [Nuñez et al., 2009, Chung and Rudgers, 2016, Lankau and Keymer, 2016] or a reduction in predator saturation [Schreiber, 2003, Gascoigne and Lipcius, 2004]), and reduced vigor owing to inbreeding and genetic drift [Fischer et al., 2000, Willi et al., 2005]. Thus, rarity may have costs that outweigh benefits.

Positive frequency-dependence also arises when a species experiences stronger negative interactions with heterospecifics than conspecifics. Mechanisms underlying PFD include niche construction via allelochemicals [Reinhart et al., 2003], changes in disturbance regime [Crandall and Knight, 2015], or nutrient cycling [Peay, 2016]. Rarity of conspecifics also increases the likelihood and costs of reproductive interference, in which mating attempts by one species have fitness costs on reproductively isolated co-occurring heterospecifics [Gröning and Hochkirch, 2008, Burdfield-Steel and Shuker, 2011, Tastard et al., 2014, Kyogoku, 2015]. Negative effects of heterospecific mating pressure can reduce use of otherwise suitable habitat [Takakura and Fujii, 2015], may be mediated by gamete interactions that reduce fertility [Hettyey et al., 2014], or even cause death. Each of these effects can impact population demography [Ting and Cutter, 2018]. The fitness costs due to reproductive interference are also often asymmetric between interacting species (see examples of magnitude, asymmetry and costs in Supplementary Table S1). PFD may, in general, be asymmetric, for example if shared mutualists provide relatively more benefit to one species than another [Sakata, 1999]. In sum, rarity is a double-edged sword, potentially providing benefits through negative frequency-dependence and competitive release, but also causing costs through diverse PFD mechanisms in both plants and animals when organisms are rare.

Environmental stochasticity may magnify the importance of PDD and PFD. If environmental fluctuations reduce population sizes of a species to the point where PDD or PFD kicks in, then such stochasticity may result in the loss of that species from the system. This phenomenon has been demonstrated in single species models with an Allee effect [Dennis, 2002, Liebhold and Bascompte, 2003, Roth and Schreiber, 2014, Schreiber, 2016]. For example, using models coupled with historical data, Liebhold and Bascompte [2003] found that environmental stochasticity could cause extinction of local gypsy moth populations (*Lymantria dispar*) in North America, even when their densities were well above the Allee threshold – the density at which the per-capita growth rate, on average, equals zero. In sharp contrast, environmental fluctuations can, via the storage effect, mediate coexistence between competing species [Chesson and Warner, 1981, Chesson, 1994]. The storage effect stabilizes coexistence, (i) when each species experiences years where environmental conditions are more favorable to it than the other species, (ii) the more common species are more limited by competition in their favorable years than the rare competitors, and (iii) species exhibit buffered growth through unfavorable years. Using data-driven models, empirical support for the storage effect exists in communities of zooplankton [Cáceres, 1997], prairie grasses [Adler et al., 2006], desert annual plants [Angert et al., 2009], tropical trees [Usinowicz et al., 2012], phytoplankton [Ellner et al., 2016], sagebrush [Chu and Adler, 2015, Ellner et al., 2016], and nectar yeasts [Letten et al., 2018]. Despite the empirical support for the storage effect and PFD, the simultaneous effects of PFD and environmental stochasticity on species coexistence is not understood. In particular, it is possible that PFD may disrupt coexistence mechanisms, like the storage effect, which rely on species having the advantage when rare.

Here, we use models to explore how positive frequency-dependence (PFD), environmental stochasticity, and asymmetry in PFD interact to influence the coexistence of species. Previous theoretical studies have considered species coexistence with resource competition and PFD by numerical simulations [Waser, 1978, Ribeiro and Spielman, 1986, Feng et al., 1997, Ruokolainen and Hanski, 2016, Molofsky et al., 2001] as well as with graphical approaches [Levin and Anderson, 1970, Kuno, 1992, Yoshimura and Clark, 1994, Kishi and Nakazawa, 2013, Kyogoku and Sota, 2017]. Although these studies revealed alternative stable states arising due to PFD, they are not well integrated into the framework of modern coexistence theory [Chesson, 2000]. To facilitate this integration, we formulated a new discrete-time model accounting for the interactive effects of competition and PFD on individual fitness. This model builds on a model that has been used extensively to empirically test and further develop coexistence theory [Adler et al., 2007, Levine and HilleRisLambers, 2009, Godoy et al., 2014, Hart et al., 2016, Godoy et al., 2017]. We present an analysis of the deterministic and stochastic versions of the model to address the following questions: How strong does niche differentiation have to be in the face of PFD to generate negative frequency-dependence and allow for coexistence? How do asymmetries in PFD and fecundity differences influence whether coexistence occurs, and can asymmetries in PFD result in non-additive effects of niche differences and PFD on coexistence? How robust is species coexistence to environmental fluctuations? How does this robustness depend on the degree of correlation between the species demographic responses to these fluctuations and temporal autocorrelations in these fluctuations? What role does the storage effect play in maintaining coexistence in the face of PFD and environmental stochasticity?

## Model and Methods

To integrate the dynamics of competition and PFD, we build on the Leslie-Gower model of competing species [Leslie and Gower, 1958] which has been used extensively for describing the dynamics of competing annual plants and insects [Leslie and Gower, 1958, Chesson, 1994, Adler et al., 2007, Godoy and Levine, 2014, Godoy et al., 2014]. The dynamics of these models are fully characterized and serve as discrete-time analogs of the classical, continuous-time Lotka-Volterra competition models [Cushing et al., 2004]. Unlike earlier models accounting for PFD [Kuno, 1992, Yoshimura and Clark, 1994, Kishi and Nakazawa, 2013], this model choice allows us to directly account for the interactive effects of PFD and competition on the ecological dynamics.

### The model

The model has two competing species with densities *N*_1_ and *N*_2_. The maximal number of offspring produced by an individual of species *i* is maximal per-capita fecundity *λ_i_*. Intra-and inter-specific competition reduce this fecundity by a linear function of the species densities. That is, let *α_ii_* and *α_ij_* be the strengths of intra- and inter-specific competition for species *i*, respectively. Then the expected number of offspring produced by an individual of species *i* experiencing no PFD is

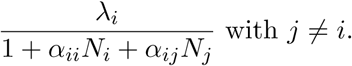

PFD independently reduces the fitness of an individual by a frequency-dependent factor

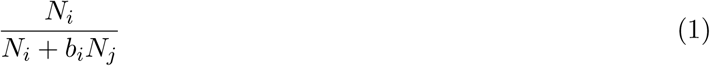
where *b_i_* determines the negative impact of species *j* on species *i*. One mechanistic interpretation of expression (1) can be given in terms of reproductive interference for species whose fecundity is more resource-limited than limited by processes involved in mating, as is found in many plants (e.g., Ghyselen et al. [2016]) and animals (e.g., Gittleman and Thompson [1988]). For such species, expression (1) can be interpreted as the probability of a successful conspecific mating.

Multiplying these components of the per-capita growth rate together yields the following deterministic model:

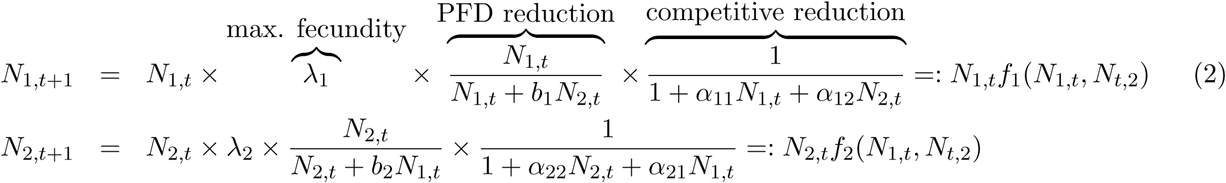
where *N_i_*,*_t_* denotes the density of species *i* in year *t*. To account for environmental stochasticity, we replace the per-capita maximal fecundities *λ_i_* with random terms *λ_i_*,*_t_* that are log-normally distributed with log means *µ_i_*, log variances 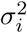, log cross-correlation *r*, and log temporal autocorrelation *τ*. The correlation *r* determines to what extent the fecundities of the two species respond in a similar manner to the environmental fluctuations. For *r* = 1, the species respond identically to the fluctuations. For *r* = 0, their responses are uncorrelated, while for *r* = −1, good years from one species are bad years for the other species. The temporal autocorrelation *τ* determines whether favorable years for one species tend to be followed by favorable years for that species (i.e., *τ* ≈ 1) or are uncorrelated to the environmental conditions in the next year (i.e., *τ* ≈ 0). As in the face of environmental stochasticity, the storage effect can promote coexistence of competing species [Chesson and Warner, 1981, Chesson, 1994], we also modify the model to account for overlapping generations that ensures population buffering, a necessary component of the storage effect. Specifically, we add terms, +*s_i_N_i_*,*_t_*, to the right hand sides of equation (2) where *s_i_* is the survival probability of an individual of species *i*.

## Methods

Our results for the deterministic model focus on highly fecund species (i.e., *λ_i_* ≫ 1) for which the analysis is substantially simpler yet still captures the full dynamical complexity of the model. Specifically, in Appendix S2, we show that the highly fecund model reduces to a one-dimensional system. For this reduced model, we find implicit expressions for the equilibria, identify their stability, and classify the dynamics into two types: contingent exclusion in which there is one unstable coexistence equilibrium and two stable single species equilibria, and contingent coexistence in which there is one stable coexistence equilibrium, two unstable coexistence equilibria, and two stable single species equilibria. To extend our analysis to the full model, we use the theory of monotone maps [Smith, 1998, Hirsch and Smith, 2005] and show, as in the high-fecundity case, that the dynamics always converge to a one-dimensional system whose dynamics are governed by a finite number of equilibria. We use this analysis and numerical computations to explore the structure of these equilibria and to determine how niche overlap, fecundity differences, and PFD interact to determine whether the models exhibit contingent coexistence or exclusion.

For the stochastic model, Appendix S3 uses results of Roth and Schreiber [2014] to show that stochastic fluctuations ultimately result in species loss whenever the species responses to the environmental fluctuations are not perfectly correlated. When species loss occurs, it occurs asymptotically in time at a super-exponential rate i.e., *N_i_*(*t*) ≈ *C* exp(− exp(*rt*)) for some *r*,*C* > 0 and for large *t*. To estimate extinction risk, we used a quasi-extinction threshold of 0.001 i.e., when a species’ density falls below 0.001, the species is declared extinct. This extinction threshold is approximately 100, 000 times smaller than the typical species density at equilibrium for the corresponding deterministic model. Our results were not qualitatively sensitive to the value of this quasi-extinction threshold. We numerically explore how the probability of species loss over finite time intervals depends on PFD, the standard deviations *σ_i_* of the environmental fluctuations, and the interspecific correlation *r* and temporal autocorrelation *τ* in these fluctuations.

To understand the role of the storage effect, we allowed for overlapping generations (i.e., positive survivorship terms *s_i_* > 0) and positive temporal autocorrelations (*τ* > 0) in the log per-capita maximal fecundities *λ_i_*. The overlapping generations ensure that the populations are buffered and the temporal autcorrelations ensure there is a positive correlation between the log per-capita fecundities and the strength of competition. These features, which are required for a storage effect [Ellner et al., 2016], are verified in Appendix S3. For this model, we computed the competitive component of the invasion growth rate of each species [Chesson, 1994, Schreiber, 2012]. Specifically, for species 2, we simulated the dynamics of species 1, in the absence of species 2, for *T* = 10, 000 time steps, to estimate its stationary distribution 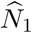. Then we estimated the invasion growth rate of species 2 without the PFD term as

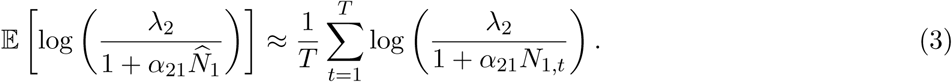

The larger the value of (3), the more quickly the species 2 would increase in the absence of positive frequency-dependence. Analytical details are provided in Appendix S3.

## Results

### Frequency-dependence, coexistence, and exclusion

Our analysis begins with the deterministic model. To ensure each species *i* can persist in isolation, we assume that the maximal fecundity *λ_i_* is greater than one for each species. Under this assumption, species *i* in isolation converges to the positive equilibrium, the carrying capacity 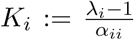. When there is PFD (*b_j_* > 0), the low density per-capita growth rate of species *j* ≠ *i* is zero at this equilibrium as individuals fail to reproduce (e.g., failure to reproduce with conspecifics due to their low frequency). Consequently, species *j* is excluded whenever it reaches such low frequencies, and the equilibria (*N*_1_,*N*_2_) = (*K*_1_, 0) and (0,*K*_2_) are locally stable.

Despite these stable, single-species equilibria, coexistence may occur at another stable equilibrium. To see when this contingent coexistence occurs, we focus on the case of highly fecund species (i.e., *λ_i_* ≫ 1) for which competition is more likely to be severe, and present analysis of the general case in Appendix S2. In this case, competitive outcomes depend on the relative per-capita growth rate (*R*_1_ : = *f*_1_*/f*_2_) of species 1 as a function of its frequency 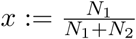. The relative per-capita growth rate of species 1 is a product of three terms (see equation (S2.3) in Appendix S2):

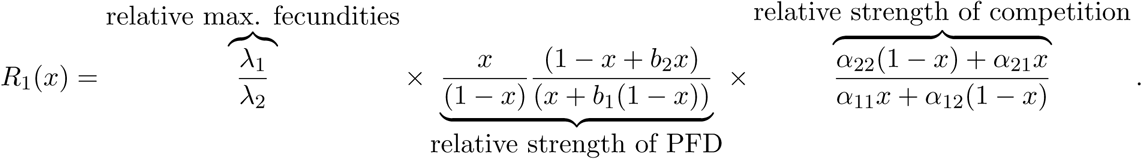

The second term increases with the frequency of species 1 whenever *b_i_* > 0 (dotted curves in Fig. 1A,C; Appendix S2).

Frequency-dependence in the third term, the relative strength of competition, can be positive or negative. The sign of this frequency-dependence depends on the niche overlap of these two species:

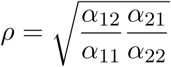

[Chesson, 2013, Godoy and Levine, 2014]. If there is partial niche overlap (*ρ* < 1), the relative per-capita growth rates without PFD exhibits negative frequency-dependence (dashed curves in Fig. 1A,C; Appendix S2). Intuitively, as a species becomes more frequent in the community, it experiences more intraspecific competition than interspecific competition; as intraspecific competition is stronger than interspecific competition, the per-capita growth rate without PFD decreases. When there is perfect niche overlap (*ρ* = 1), the relative per-capita growth rate without PFD is frequency-independent. In this case, the relative per-capita growth rate only exhibits PFD and coexistence is not possible. Consequently, from now on, we assume *ρ* < 1.

**Figure 1:**
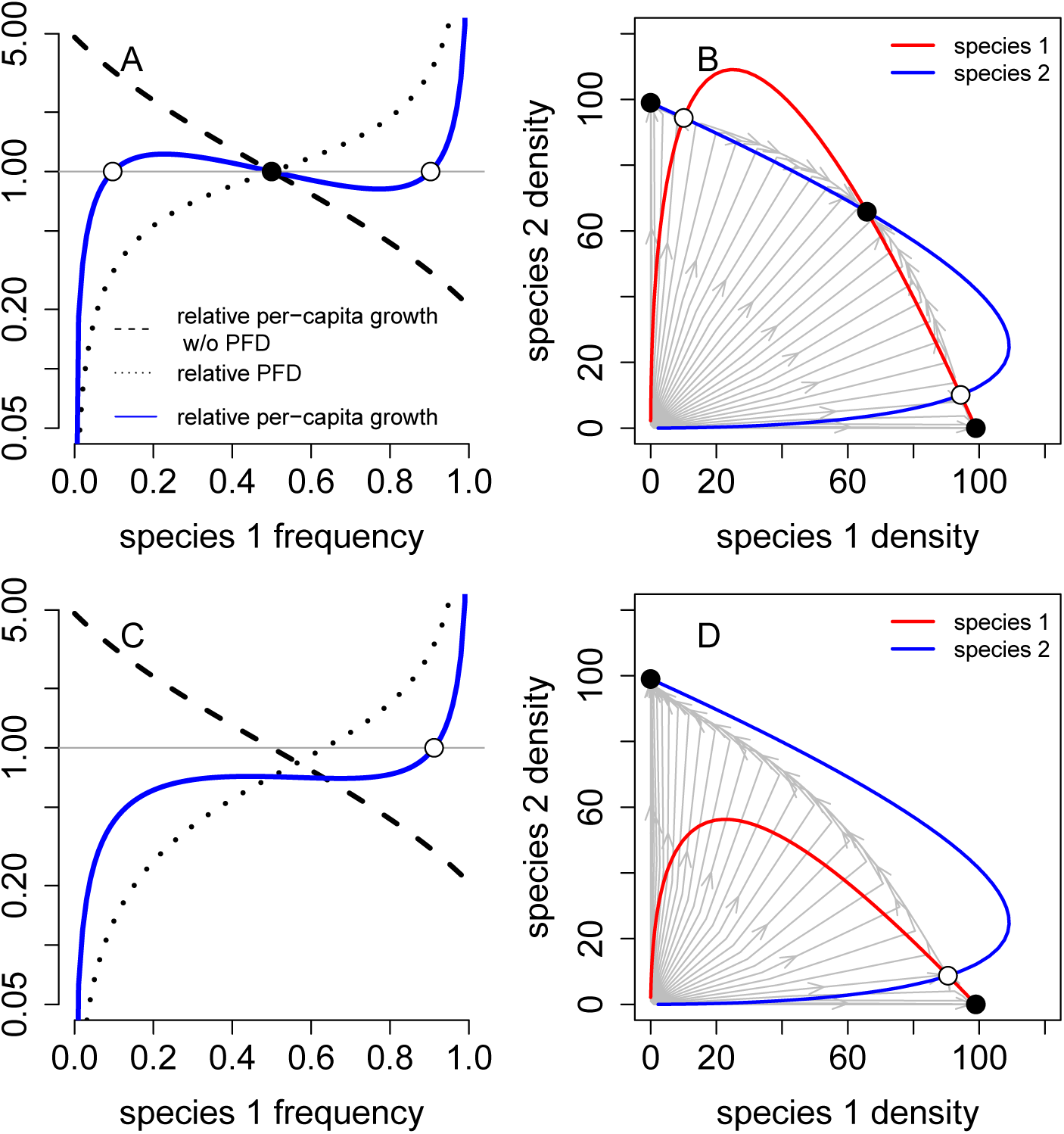
Frequency-dependent feedbacks and the dynamics of contingent coexistence (A,B) and contingent exclusion (C,D). In A and C, relative strength of PFD (black dotted), relative strength of competition (black dashed), and relative per-capita capita growth rate *R*_1_(*x*) (blue) for species 1 are shown as a function of the frequency *x* of species 1. In B and D, colored curves correspond to the zero-growth nullclines, and trajectories for different initial conditions are gray lines. In all figures, stable equilibria/frequencies are filled circles and unstable equilibria/frequencies are unfilled circles. In A and B, low niche overlap results in negative frequency-dependence at intermediate species frequencies and coexistence. In C and D, large niche overlap result in PFD in the relative per-capita growth rate of species 1 at all species 1’s frequencies. Parameter values are *b*_1_ = *b*_2_ = 0.25 in A and B, and *b*_1_ = 0.75 and *b*_2_ = 0.25 in C and D. Other parameter values are *λ_i_* = 100, *α_ii_* = 1, and *α_ij_* = 0.2.

Provided there is sufficiently low niche overlap, the relative per-capita growth rate *R*_1_ of species 1 exhibits negative frequency-dependence at intermediate species frequencies (Fig. 1A,B). When this occurs, there are two critical frequencies, *x*_low_ < *x*_high_ of species 1 such that (i) the per-capita growth rate of species 1 is greater than the per-capita growth rate of species 2 when its frequency is slightly above *x*_low_, and (ii) the per-capita growth rate of species 2 is greater than the per-capita growth rate of species 1 when species 1’s frequency is slightly below *x*_high_. When species 1’s frequency lies between *x*_low_ and *x*_high_, negative frequency-dependent feedbacks dominate and the species approach a unique stable coexistence equilibrium. In contrast, when species 1’s frequency falls below *x*_low_ or exceeds *x*_high_, PFD feedbacks dominate and either species 1 gets excluded by species 2 or excludes species 2, respectively.

When niche overlap is too great, PFD dominates at all species frequencies and coexistence is not possible (Fig. 1C,D). Consequently, there is a critical frequency *x*_bistable_ of species 1 below which species 1 is excluded and above which species 2 is excluded.

### Niche overlap, fecundity differences, and contingent coexistence

To better understand when coexistence or exclusion occurs, we focus on the case where the species are demographically similar with respect to competition (*α*_11_ = *α*_22_ and *α*_12_ = *α*_21_) but potentially differ in their maximal fecundities (*λ_i_*) or their susceptibility to PFD (*b_i_*). The general case is presented in Appendix S2. If there is no PFD (*b*_1_ = *b*_2_ = 0), coexistence occurs if the niche overlap is less than the the ratio of maximal fecundities *λ_i_/λ_j_*:

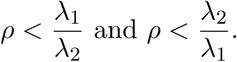

In this case, coexistence is not contingent upon initial conditions. This coexistence condition is sharp: if it is satisfied, the species coexist, else they do not (Fig. 2A).

When species experience both positive frequency-dependence as well as negative frequency-dependence due to interspecific competition, coexistence requires that the additive effects of niche overlap and the strength of PFD are less than the ratio of maximal fecundities:

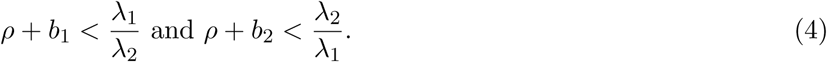

If (4) is not satisfied, negative frequency-dependent feedbacks are too weak to promote coexistence. Satisfying (4), however, need not ensure coexistence due to nonlinear, interactive effects between PFD and niche overlap (the distance between the dashed lines and the coexistence regions in Fig. 2B-D). Equation (S2.5) in Appendix S2 provides an explicit condition for this nonlinear interactive. In general, this expression is difficult to interpreet biologically. However, for species that exhibit no differences in maximal fecundities (*λ*_1_ = *λ*_2_) and are equally susceptible to PFD (*b*_1_ = *b*_2_ = *b*), this interactive effect equals 3*ρb* and coexistence occurs if (Appendix S2)

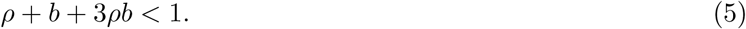

As niche overlap and PFD contribute equally to this nonlinear interactive effect, coexistence is least likely when the strength of PFD and niche overlap are equally strong (Fig. 2B).

Differences in the maximal fecundities or asymmetries in the strength of PFD lead to larger, nonlinear effects on coexistence (the greater distance between the dashed line and the coexistence region in Figs. 2C,D than B). When PFD is symmetric, larger differences in the maximal fecundities (e.g., larger values of *λ*_1_*/λ*_2_) always inhibit coexistence (Fig. 2C). Numerical simulations suggest that interactive effects of PFD and niche overlap continue to be symmetric in this case. When differences in the maximal fecundities are too large to permit coexistence, the species with the fecundity disadvantage can be excluded despite being at an initially higher frequency.

**Figure 2:**
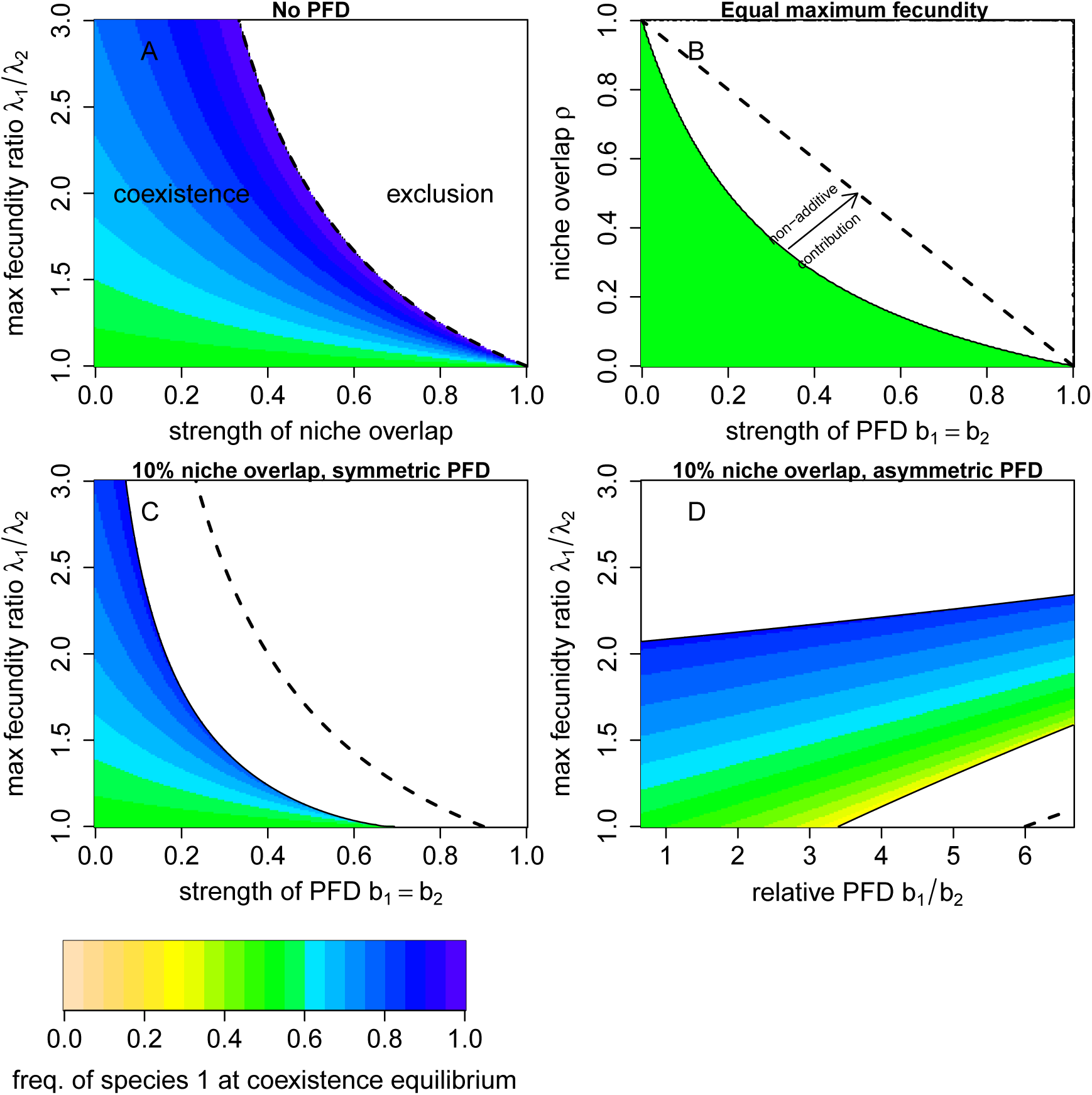
Effects of PFD, niche overlap, and fecundity differences on species coexistence. Colored region corresponds to contingent coexistence, while the white region corresponds to exclusionary dynamics. Colored shading indicate frequency of species 1 at the stable coexistence equilibrium. In A, B, and C, species are equally sensitive to PFD (*b*_1_ = *b*_2_) or niche overlap (*α*_12_ = *α*_21_). In A, there is no PFD (*b_i_* = 0), while both PFD and niche overlap occur in B (*α_ij_* > 0 and *b_i_* > 0). In C and D, species differ in their maximal fecundities and experience either symmetric (in C) or asymmetric (in D) PFD. Dashed lines the coexistence boundary from equation (4) which only accounts for the additive contributions of PFD and niche overlap. The solid black lines are determined by the analytic criterion presented in Appendix S2. In C and D, there is symmetric 10% niche overlap (*α_ij_* = 0.1). Other parameter values are *λ*_2_ = 100 and *α_ii_* = 1.

When there are sufficiently strong asymmetries in PFD (*b*_1_*/b*_2_ > 3.5 in Fig. 2D), coexistence occurs at intermediate differences in the maximal fecundities. If the fecundity advantage of the species 1 is not sufficiently high, coexistence is not possible and this species can be excluded even when it is initially at the higher frequency (Fig. 2D). Alternatively, if the fecundity advantage of species 1 is too large, coexistence is not possible and species 2 has a lower threshold frequency below which it is excluded (Fig. 2D).

**Figure 3:**
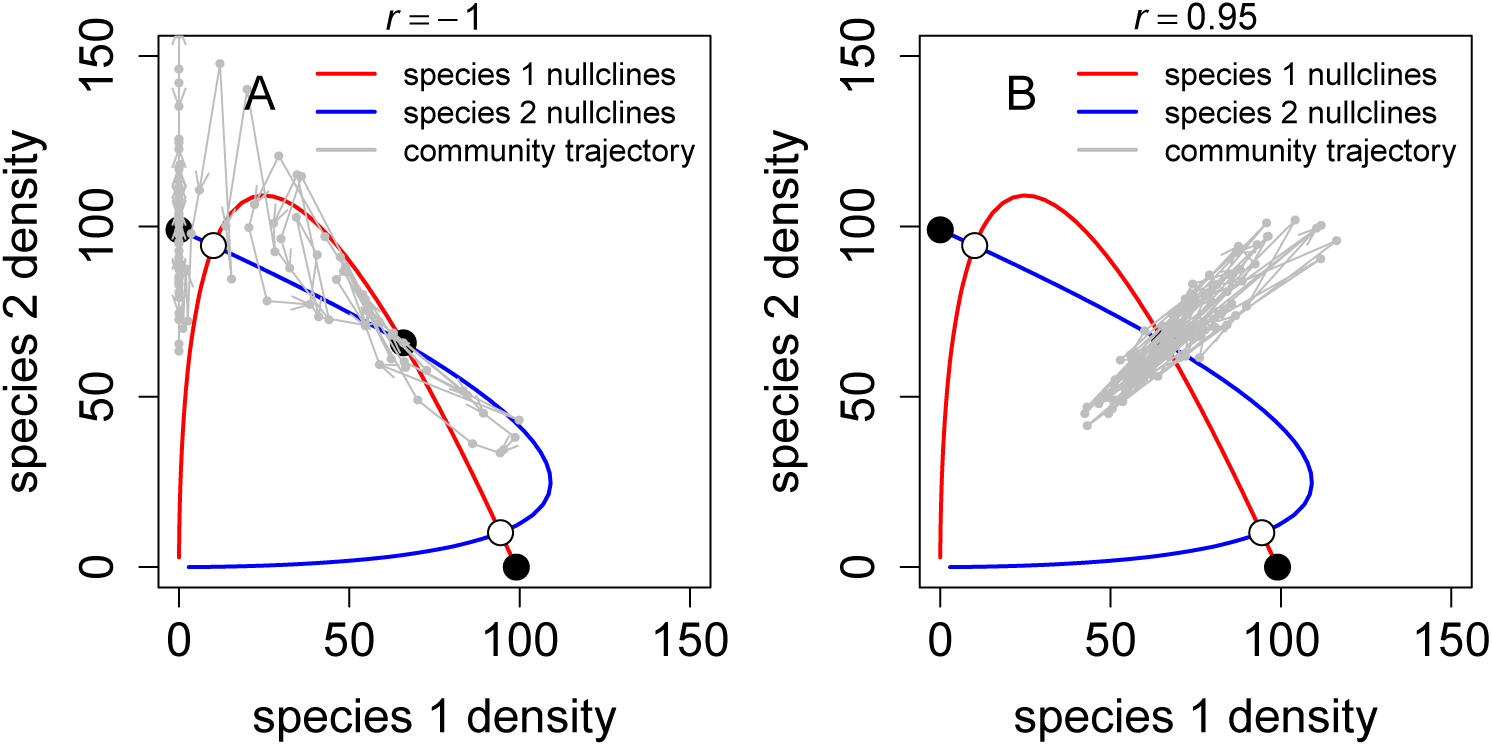
Fluctuations, correlations, and coexistence. In (A) and (B), 100 year simulations (gray lines) of the stochastic model with fluctuating maximal fecundities are plotted in the phase plane. The nullclines for the mean field model shown as red and blue curves, and the corresponding stable and unstable equilibria for the mean field model as solid and unfilled circles, respectively. In (A) and (B), the correlations *r* between the log maximal fecundities of the species are negative (*r* = −1) and positive (*r* = 0.95), respectively. Parameter values as in Figure 1 with 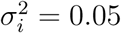 and *τ* = 0.

### Stochastic environments

When the maximal fecundities *λ_i_*,*_t_* fluctuate stochastically, the fluctuations in the frequency dynamics are determined by the fluctuations in the ratio of these fecundities *λ*_1,_*_t_/λ*_2,_*_t_*. As these fecundities *λ_i_*,*_t_* are log-normally distributed with log-mean *µ_i_*, log-variance 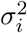 and correlation *r*, their ratio *λ*_1,_*_t_/λ*_2,_*_t_* is log-normally distributed with

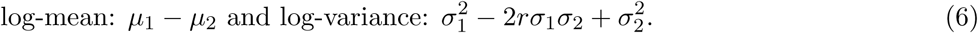

Equation 6 implies that positively correlated responses (*r* > 0) of the two species to the environmental fluctuations decrease the log-variance in the frequency dynamics. Intuitively, environmental fluctuations cause the fecundities of each species to change by the same factor and, thereby, reduces the effect of these fluctuations on the ratio of maximal fecundities (Fig. 3B). Indeed, when the responses to environmental fluctuations are of the same magnitude and perfectly correlated (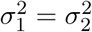 and *r* = 1), there are no fluctuations in these fecundity ratios and species may coexist indefinitely.

In contrast, when species exhibit opposing responses to environmental fluctuations (*r* < 0), environmental fluctuations that drive one species to higher densities simultaneously drives the other species to low densities. This behavior results in larger fluctuations in the species frequencies (Fig. 3A). In the extreme case where the responses to the environmental fluctuations are of the same magnitude and are perfectly negatively correlated (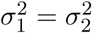 and *r* = −1), the fluctuations in the log ratio of fecundities are twice as large as that for uncorrelated fluctuations (i.e., 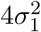 versus 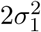).

**Figure 4:**
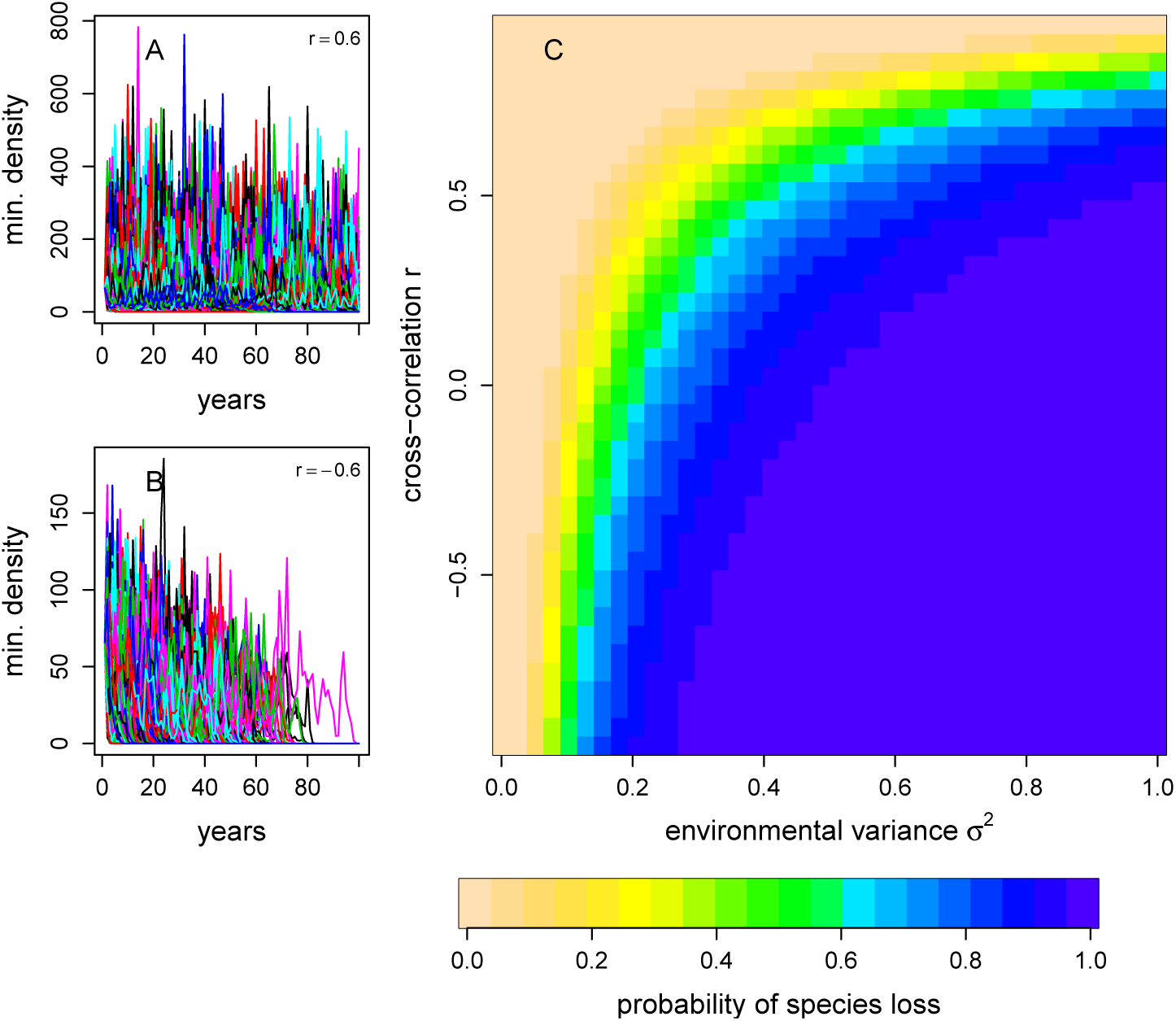
Extinction increases with variation in environmental fluctuations and decreases with correlated species responses to the environment. In the left panels, plots of the minimum of the species densities (i.e., the smaller value of N_1,_*_t_* and *N*_2,_*_t_*) for multiple simulations of the stochastic model. In (A), the species have negatively correlated responses to the environment (cross-correlation coefficient *r* < 0). In (B), the species positively correlated responses to the environment (*r* > 0). In the right panel (C), a contour plot of the probability of the loss of a species within 50 years with respect to the magnitude of the fluctuation *σ*_1_ = *σ*_2_ = *σ* and the cross-correlation *r*.

When species exhibit some differentiated responses to environmental fluctuations (*r* < 1 or 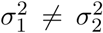), environmental fluctuations ultimately will drive one of the species extinct, whether or not deterministic coexistence is possible (Appendix S3, Fig. 4). Intuitively, environmental fluctuations can push one of the species to a sufficiently low frequency that the deterministic effects of PFD rapidly drive the species to extinction. Larger environmental fluctuations increase the likelihood of these events and, thereby, increase the probability of species loss. As negative correlations increase fluctuations in frequencies, they also increase the likelihood that one species falls below its critical frequency and rapidly goes extinct. Consequently, the probability of extinction decreases with positive correlations. In fact, when species responses to the environmental fluctuations are identical (i.e., 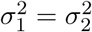 and *r* = 1), environmental stochasticity does not drive any species extinct provided they are initially near a stable, coexistence equilibrium of the deterministic model (Appendix S3). In contrast to the effects of positive cross-correlations, positive temporal autocorrelations increase quasi-extinction risk (Fig. S3-1). Intuitively, positive temporal autocorrelation leads to longer runs of unfavorable conditions to one species thereby making this species more vulnerable to PFD.

**Figure 5:**
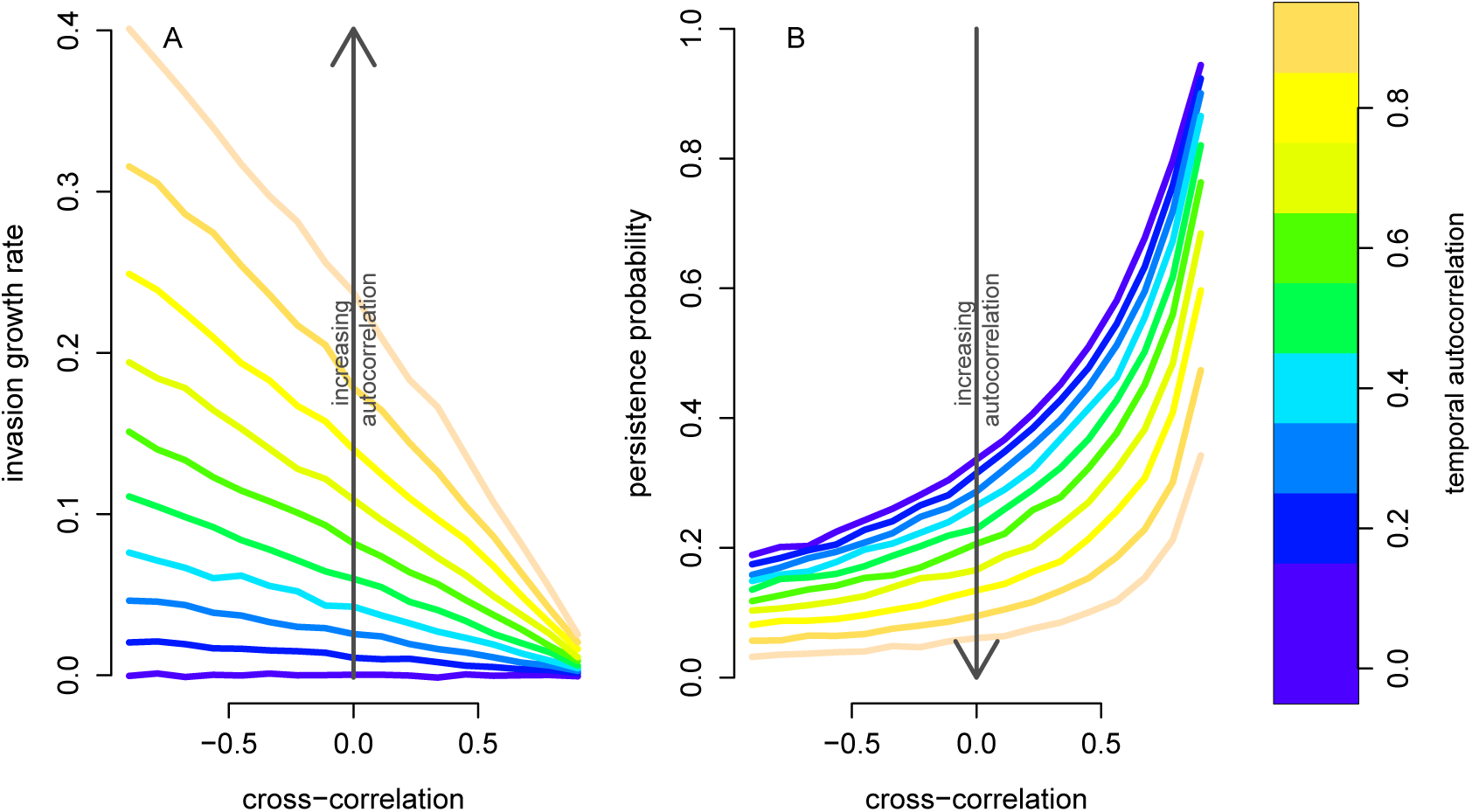
The storage effect does not overcome the effects of PFD. The competitive component of the invasion growth rate (A) that measures the strength of the storage effect, and the probability of extinction (B) in 50 years are shown for different values of the cross-correlation *r* and temporal-autocorrelation. Parameter values: *α*_11_ = *α*_12_ = *α*_21_ = *α*_22_ = 1, *λ*_1_ = *λ*_2_ = 100, *b*_1_ = *b*_2_ = 0.01, *s*_1_ = *s*_2_ = 0.5.

To understand whether the storage effect could alter the predictions about the effects of cross-correlations on extinction risk, we simulated the model with overlapping generations (i.e., *s_i_* > 0) and varying levels of temporal autocorrelation *τ*. To strengthen the potential for a storage effect [Chesson, 1994], species had no difference in their mean maximal fecundities, exhibited complete niche overlap (i.e., *α*_11_ = *α*_22_ = *α*_12_ = *α*_21_), experienced weak positive frequency-dependence (i.e., *b*_1_ = *b*_2_ = 0.01), and had high survival (i.e., *s* = 0.5). The competitive component of the invasion growth rate (see definition in *Methods*) when rare decreased with positive cross-correlations and increased with positive temporal autocorrelations (Fig. 5A). Therefore, the storage effect is greatest with negative cross-correlations and positive temporal autocorrelations. Despite this trend, the probability of extinction increased with negative cross-correlations and positive temporal autocorrelations in the maximal fecundity values (Fig. 5B). Namely, the impact of these correlations on extinction risk by generating low frequencies of one species outweighed their impact of increasing the long-term per-capita growth rates when rare.

The effects of asymmetries in maximal fecundities and PFD on extinction risk largely follow patterns suggested by the deterministic model: when a species is at low frequency at the stable, coexistence equilibrium, extinction risk is greater (compare Fig. 2D to Fig. 6). In particular, for a given level of asymmetry in PFD, persistence of both species is most likely at an intermediate fecundity advantage of the species more susceptible to PFD. For smaller differences in maximal fecundity, the species with the fecundity advantage is more likely to go extinct. For larger fecundity differences, the species less susceptible to PFD is more likely to go extinct. As larger fecundity differences (i.e., *µ*_1_ larger than *µ*_2_) result in larger fluctuations in their ratio (i.e., variance *λ*_1,_*_t_/λ*_2,_*_t_* equals 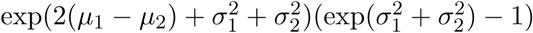 when *r* = 0), extinction risk is generally greater due to larger fecundity differences rather than smaller fecundity differences (blue region larger in Fig. 6B than in Fig. 6A).

**Figure 6:**
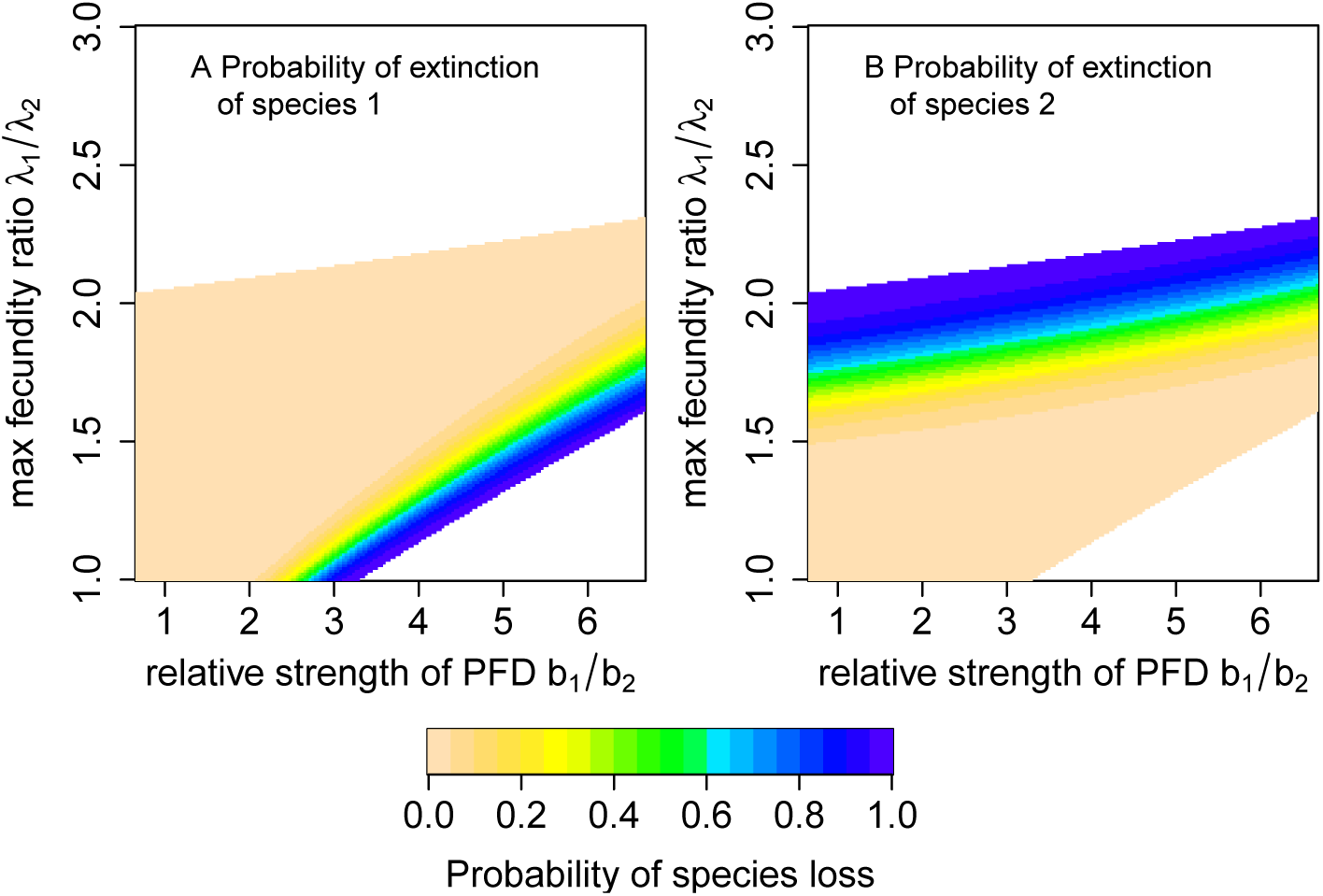
Fluctuations, asymmetric PFD, and extinction risk. The probability of extinction of species 1 (A) or species 2 (B) within 100 years. Species started at the stable coexistence equilibrium for the mean field model (i.e., deterministic model with 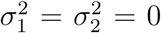) whenever it exists. Parameter values: 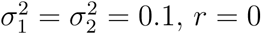, and remaining parameters as in Fig. 2D.

## Discussion

Many competing species are likely to experience both negative and positive frequency-dependence. Positive frequency-dependence (PFD), in and of itself, does not allow for coexistence and leads to alternative stable states supporting only a single species [Amarasekare, 2002, Fukami and Nakajima, 2011]. In contrast, negative frequency-dependence allows for stable coexistence [Adler et al., 2007] but does not allow for alternative stable states. For competing species experiencing both positive and negative frequency-dependent feedbacks, a new dynamic emerges supporting alternative stable states including ones at which the species coexist (Fig. 1). This dynamic can occur when positive frequency-dependence occurs at low species frequencies, and negative frequency-dependence dominates at intermediate species frequencies. More generally, this dynamic arises when there are multiple changes in the sign (i.e., positive versus negative) of frequency dependence. When these conditions are met, we find that coexistence is determined by more complex interactions of both positive and negative frequency-dependence, rather than the “mutual invasibility criterion” of modern coexistence theory [Chesson, 2000] or by species growth rates when rare [Hofbauer and Sigmund, 1998, Schreiber, 2000, Schreiber et al., 2011]. Our deterministic analysis highlights that niche overlap and PFD have negative synergistic effects on coexistence, yet trade-offs between PFD and fecundity can facilitate deterministic coexistence.

Deterministic coexistence requires that the additive effects of niche overlap and PFD need to be smaller than the ratio of maximal fecundities (see equation (4)). When both competitors experience the same strength of PFD, the interactive effects of PFD and niche overlap are symmetric: a simultaneous increase in the strengths of PFD and niche overlap have a more negative impact on coexistence than increasing the strength of one more than the other (see equation (5)). Asymmetries in the strength of PFD, which are common (e.g., Supplementary Table S1), can facilitate coexistence if the species more vulnerable to PFD has the higher maximal fecundity. This trade-off is affected by niche overlap as highlighted in the coexistence condition (4). For example, a twofold advantage in fecundity for one species requires that the other species’ vulnerability to PFD must be more than 50% less for coexistence (i.e., if *λ*_1_ = 2*λ*_2_, then b_2_ < 1/2 − *ρ* whereas *b*_1_ < 2 − *ρ*). The greater niche overlap, the stronger the trade-off needs to be. Given the empirical prevalence of asymmetries in PFD for coexisting competitors (e.g., the asymmetries in reproductive interference reported in Table S1), our results suggest there may be a countervailing strong trade-off in the maximal fecundities or sensitivities to competition (see below) for these species pairs.

Our deterministic analysis complements and extends earlier work by Kishi and Nakazawa [2013], who analyzed the dynamics of two competing species experiencing reproductive interference. Unlike our discrete-time model, which accounts for the simultaneous effects of PFD and competition on fecundity, Kishi and Nakazawa [2013]’s model is continuous-time and assumes that competition increases mortality rates while reproductive interference reduces birth rates. Thus, our model integrates more naturally into the framework of modern coexistence theory [Chesson, 2000, Adler et al., 2007] and is readily applicable to PFD in annual plants [Levine and HilleRisLambers, 2009, Godoy and Levine, 2014, Hart et al., 2016] and insects (see also Ribeiro and Spielman [1986]). Under the assumption of symmetry in reproductive interference, Kishi and Nakazawa [2013] derived a similar coexistence condition to our condition (4) that holds for asymmetric, as well as symmetric, PFD. Kishi and Nakazawa [2013] numerically demonstrated trade-offs between the strength of reproductive interference and sensitivity to competition, which determines niche overlap, could facilitate coexistence; a finding that complements our results about trade-offs between sensitivity to PFD and maximal fecundity.

Just as strong positive density-dependence can make populations particularly vulnerable to environmental fluctuations [Courchamp et al., 1999, Dennis, 2002, Liebhold and Bascompte, 2003, Roth and Schreiber, 2014], environmental fluctuations make coexistence of competitors experiencing PFD more tenuous. Indeed, our analysis reveals that asymptotic extinction is inevitable when either of the competitors are repeatedly pushed over the critical threshold where positive frequency-dependence kicks in. Despite this extinction only occurring asymptotically, the rare species decreases at a super-exponential rate (i.e., the species’ population size decays faster than *e^rt^* for any *r* < 0) and, thus, would rapidly go extinct due to demographic stochasticity. This extinction risk is most severe for interacting species exhibiting opposing demographic responses to environmental fluctuations e.g., one species producing more offspring in cooler years while the other species produces more offspring in warmer years. Specifically, these negatively correlated responses result in greater fluctuations in the relative frequencies of the species and, therefore, are more likely to drive one of them to sufficiently low frequencies, at which point PFD kicks in. In sharp contrast, for species with highly positively correlated responses to environmental fluctuations, fluctuations in species frequencies are minimal and extinction risk is much smaller.

These effects of species’ correlated responses to environmental conditions are in direct opposition to Chesson’s storage effect [Chesson and Warner, 1981, Chesson, 1994]. The storage effect promotes coexistence when there are (i) species-specific responses to environmental conditions, (ii) covariance between environmental conditions and the strength of competition, and (iii) buffered population growth. The first ingredient (i) is strongest when the species exhibit negatively correlated responses to the environment and the weakest when the species exhibit nearly perfectly correlated response to the environment. The second ingredient is strongest when there are positive temporal autocorrelations and weakest when there are temporally uncorrelated fluctuations. Our analysis reveals that even when the storage effect is operating, the effect of PFD on coexistence due to correlated responses to the environment and temporal autocorrelations outweigh the opposing effects of the storage effect. In particular, even though negatively correlated responses to the environment increase the strength of the storage effect, the increased variation in relative frequencies of the species leads to greater extinction risk. The reason for this is two-fold. First, when one species becomes rare, its reduction in fitness due to PFD is sufficiently strong to eliminate any signature of the storage effect. Thus, the storage effect can only operate when neither species is too rare. However, at these more intermediate frequencies, even the less common species is experiencing more intraspecific competition which, in and of itself, dilutes the strength of the storage effect. Hence, our results predict that coexisting species simultaneously exhibiting the storage effect and PFD should be uncommon.

While our theory provides a first step in developing a community ecology theory accounting for positive frequency-dependence and environmental fluctuations, there are many additional complexities that need to be explored. These complexities include interactions between PFD and spatial population structure [Ruokolainen and Hanski, 2016], interference competition [Amarasekare, 2002], evolution toward avoiding PFD (i.e., reproductive character displacement or reproductive interference-driven niche partitioning [Liou and Price, 1994, Goldberg and Lande, 2006]), and conservation of rare species by considering the interaction between genetic and demographic swamping [Todesco et al., 2016]. For example, aggregative behavior of species may allow species at low frequency in the larger community to be partially buffered from positive frequency-dependent processes by creating tiny local patches of higher density [Molofsky et al., 2001, Ruokolainen and Hanski, 2016]. Developing a theory to understand how these many forms of positive frequency-dependence interact with environmental fluctuations to determine community structure is a major challenge that will likely require a paradigm shift in coexistence theory.

## Acknowledgments

We would like to thank two anonymous reviewers and R. Iritani for their extensive and thoughtful comments. SJS was supported by the US National Science Foundation Grants DMS1313418,1716803. MY was supported by the Japan Society for the Promotion of Science (JSPS) Grant-in-Aid for Scientific Research (KAKENHI) 15H02642, 16K18618, and 16H04846 and by John Mung Program and Hakubi Center for Advanced Research of Kyoto University. SYS acknowledges NSF DEB 13-42841 and the California Agricultural Experiment Station.

## Appendix S2 Deterministic Analysis

### Analysis of the case of symmetric species

When the two species are identical (*b* = *b*_1_ = *b*_2_, *λ* = *λ*_1_ = *λ*_2_, *α* = *α*_11_ = *α*_22_, and *β* = *α*_12_ = *α*_21_), the analysis can be simplified. This symmetric version of the model has a coexistence equilibrium, 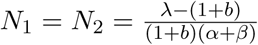. For the equilibrium to be present and locally stable, the following conditions must hold:

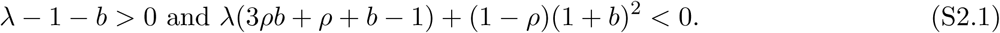
where *ρ* = *β/α* in this symmetric case. This is equivalent to the inequality (3) in Kishi and Nakazawa [2013]. Thus *λ* and *ρ* are equivalent to *b/d* and *c*, where *b*, *d*, and *c* are birth rate, death rate, and the relative strength of interspecific to intraspecific resource competition in the ODEs of Kishi and Nakazawa [2013]. This indicates that decreasing *λ* value results in reduced parameter regions for coexistence in the *b*-*β* plane (as in Fig. 2B).

### Analysis of the case of high fecundity: *λ_i_* ≫ 1

When *λ_i_* ≫ 1 for both species, we show that model (2) can be approximated by

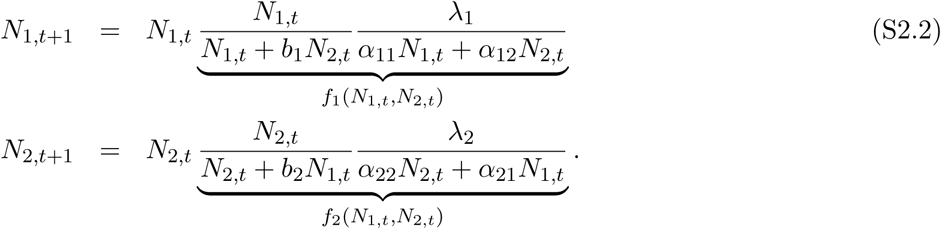

To show that the simplified model (S2.2) provides a good description of the dynamics of the global attractor of (2), we begin by showing that non-zero solutions of (S2.2) ultimate yield population sizes of order *λ_i_*. To this end, let *A* = max{*α*_11_,*α*_22_,*α*_12_,*α*_21_}, *B* = max{*b*_1_,*b*_2_, 1}, and *R* = min{*λ*_1_,*λ*_2_}. If *n_t_* = *N*_1,_*_t_* + *N*_2,_*_t_*, then

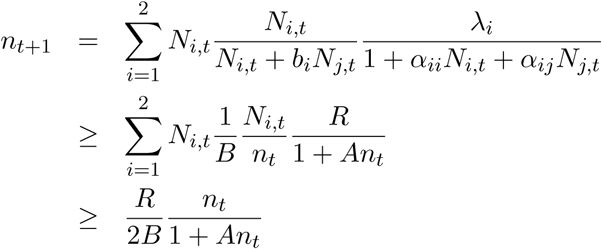
where the final inequality follows from *a*^2^ + *b*^2^ ≥ (*a* + *b*)^2^*/*2 for any *a* ≥ 0 and *b* ≥ 0. Hence, by monotonicity, if *n*_0_ > 0 and *R* ≥ 4*B*, then

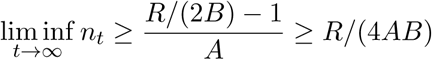

In particular, if we define *a* = min{*α*_11_,*α*_22_,*α*_12_,*α*_21_} and *R* ≥ 4*B*, then *α_ii_N_i_*,*_t_* + *α_ij_N_j_*,*_t_* ≥ *aR/*(4*AB*) for sufficiently large *t*. Hence, provided that *N*_1,0_ + *N*_2,0_ > 0 and *R* ≥ 4*B*, (*N*_1,_*_t_,N*_2,_*_t_*) enters the set

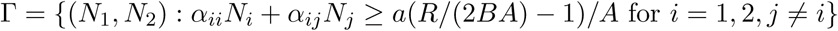
for *t* sufficiently large.

**Table S1:**
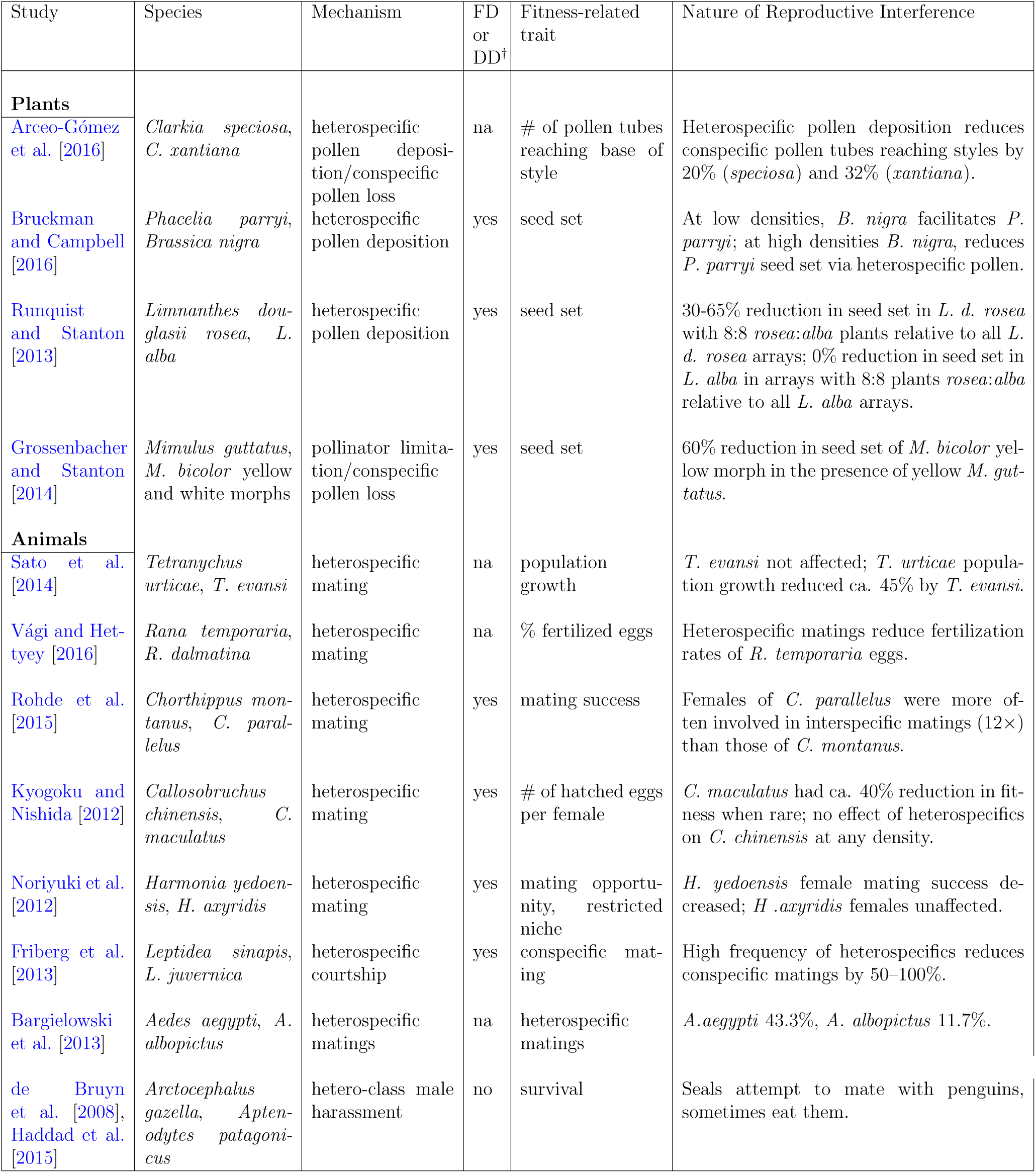
Examples of reproductive interference, a particular form of PFD, since the review of Gröning and Hochkirch [2008] from diverse plants and animals. ^†^Frequency-dependence or density-dependence.

| Study | Species | Mechanism | FD or DD<sup>†</sup> | Fitness-related trait | Nature of Reproductive Interference |
| --- | --- | --- | --- | --- | --- |
| **Plants** |  |  |  |  |  |
| Arceo-Gómez et al. [2016] | *Clarkia speciosa*, *C. xantiana* | heterospecific pollen deposition/conspecific pollen loss | na | # of pollen tubes reaching base of style | Heterospecific pollen deposition reduces conspecific pollen tubes reaching styles by 20% (*speciosa*) and 32% (*xantiana*). |
| Bruckman and Campbell [2016] | *Phacelia parryi*, *Brassica nigra* | heterospecific pollen deposition | yes | seed set | At low densities, *B. nigra* facilitates *P. parryi*; at high densities *B. nigra*, reduces *P. parryi* seed set via heterospecific pollen. |
| Runquist and Stanton [2013] | *Limnanthes douglasii rosea*, *L. alba* | heterospecific pollen deposition | yes | seed set | 30-65% reduction in seed set in *L. d. rosea* with 8:8 *rosea:alba* plants relative to all *L. d. rosea* arrays; 0% reduction in seed set in *L. alba* in arrays with 8:8 plants *rosea:alba* relative to all *L. alba* arrays. |
| Grossenbacher and Stanton [2014] | *Mimulus guttatus*, *M. bicolor* yellow and white morphs | pollinator limitation/conspecific pollen loss | yes | seed set | 60% reduction in seed set of *M. bicolor* yellow morph in the presence of yellow *M. guttatus*. |
| **Animals** |  |  |  |  |  |
| Sato et al. [2014] | *Tetranychus urticae*, *T. evansi* | heterospecific mating | na | population growth | *T. evansi* not affected; *T. urticae* population growth reduced ca. 45% by *T. evansi*. |
| Vági and Hettyey [2016] | *Rana temporaria*, *R. dalmatina* | heterospecific mating | na | % fertilized eggs | Heterospecific matings reduce fertilization rates of *R. temporaria* eggs. |
| Rohde et al. [2015] | *Chorthippus montanus*, *C. parallelus* | heterospecific mating | yes | mating success | Females of *C. parallelus* were more often involved in interspecific matings (12×) than those of *C. montanus*. |
| Kyogoku and Nishida [2012] | *Callosobruchus chinensis*, *C. maculatus* | heterospecific mating | yes | # of hatched eggs per female | *C. maculatus* had ca. 40% reduction in fitness when rare; no effect of heterospecifics on *C. chinensis* at any density. |
| Noriyuki et al. [2012] | *Harmonia yedoensis*, *H. axyridis* | heterospecific mating | yes | mating opportunity, restricted niche | *H. yedoensis* female mating success decreased; *H. axyridis* females unaffected. |
| Friberg et al. [2013] | *Leptidea sinapis*, *L. juvernica* | heterospecific courtship | yes | conspecific mating | High frequency of heterospecifics reduces conspecific matings by 50–100%. |
| Bargielowski et al. [2013] | *Aedes aegypti*, *A. albopictus* | heterospecific matings | na | heterospecific matings | *A. aegypti* 43.3%, *A. albopictus* 11.7%. |
| de Bruyn et al. [2008], Haddad et al. [2015] | *Arctocephalus gazella*, *Aptenodytes patagonicus* | hetero-class male harassment | no | survival | Seals attempt to mate with penguins, sometimes eat them. |

Now assume that *λ*_2_ = *Cλ*_1_ for some *C* > 0 and *λ*_1_ ≫ max{1, 4*B*}. As (2) is equivalent to

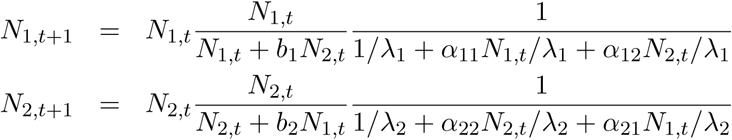

Let *c* = min{1, 1*/C*}. For (*N*_1_,*N*_2_) ∈ Γ, we have that *α_ii_N_i_/λ_i_* + *α_ij_N_j_/λ_i_* ≥ *c/*(4*BA*) > 0. Hence provided that (*N*_1_,*N*_2_) ∈ Γ and *λ*_1_ is sufficiently large

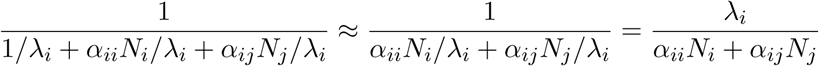
and we have justified the approximation (S2.2) of the long-term dynamics of (2).

### Analysis of the simplified model (S2.2)

Equation S2.2 can be reduced to a one-dimensional model via the change of coordinates 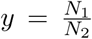. In this coordinate system

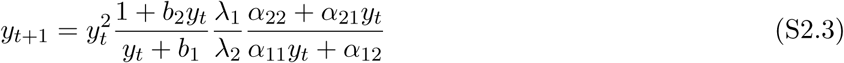

In particular, (S2.3) with *x* = *y* justifies the expression for the relative fitness *R*_1_(*x*) in the main text. For our analysis of (S2.3), we first verify the two assertions in the main text about the relative fitness function and then analyze the dynamics of (S2.3).

In the *y* coordinate system, the relative fitness function *f*_1_*/f*_2_ is given by

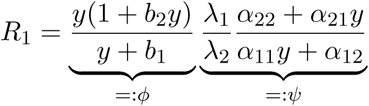
where *ϕ* corresponds to the “relative mating success” of species 1 and *ψ* corresponds to the “relative reproductive potential” of species 1. A direct computation of the derivatives yields:

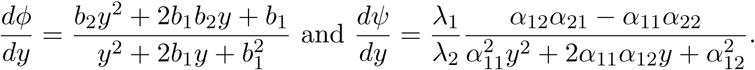
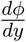 is always positive, and 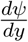 is negative if and only if *α*_11_*α*_22_ > *α*_12_*α*_21_ i.e., *ρ* < 1. To get the derivatives with respect to the frequency (*x* = *N*_1_*/*(*N*_1_ + *N*_2_)) of species 1, notice that *y* = *x/*(1 − *x*) and 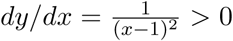 for *x* ≠ 1. Therefore, by the chain rule

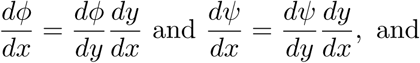
the derivatives *dϕ/dx* and *dψ/dx* have the same signs as the derivatives *dϕ/dy* and *dψ/dy*, respectively. Therefore, as claimed in the main text, *ϕ* is an increasing function of *x* and *ψ* is a decreasing function of *x* if and only if *α*_11_*α*_22_ > *α*_12_*α*_21_ i.e., *ρ* > 1.

Now we perform the dynamical analysis of (S2.3). As the right-hand side of (S2.3) is an increasing function of *y* for *x* ≥ 0, all solutions either approach an equilibrium or approach +∞ which corresponds to species 2 going extinct in (S2.2). The non-zero (and non-infinite) equilibria are given by the solutions of *R*_1_ = 1. Equivalently, the roots of the cubic:

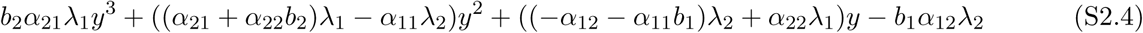

Contingent coexistence occurs when there are three positive solutions to this cubic equation i.e., two unstable equilibria and one stable equilibrium. Contingent exclusion occurs when there is only one positive real root to the equation. As this equation is negative at *y* = 0 and approaches +∞ as *y* → +∞ (i.e., the cubic coefficient is positive), there never can be exactly two positive solutions.

As the cubic (S2.4) is negative at *y* = 0 and approaches +∞ as *y* → +∞, three positive solutions are possible only if the first derivative of this cubic is positive at *y* = 0 and the second derivative at *y* = 0 is negative. Namely,

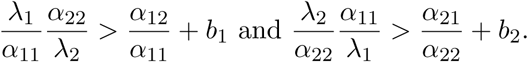

In the special case where *α*_11_ = *α*_22_ and *α*_12_ = *α*_21_, the ratios *β_i_/α_i_* equal *ρ*, and the coexistence condition becomes

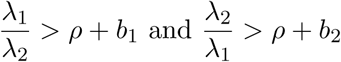
as presented in the main text.

While these coexistence conditions are only necessary, a necessary and sufficient condition is given by considering the discriminant of the (S2.4). Specifically, define

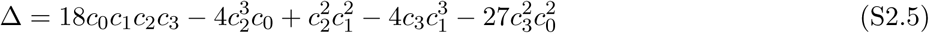
where *c*_0_ = −*b*_1_*α*_12_*λ*_2_, *c*_1_ = ((−*α*_12_ − *α*_11_*b*_1_)*λ*_2_ + *α*_22_*λ*_1_), *c*_2_ = ((*α*_21_ + *α*_22_*b*_2_)*λ*_1_ − *α*_11_*λ*_2_), and *c*_3_ = *b*_2_*α*_21_*λ*_1_. If Δ > 0, then (S2.4) has three real roots which are all positive if the necessary condition for coexistence is satisfied. Unfortunately, while this expression provides a quick computational method for identifying whether coexistence occurs or not, it is not readily interpretable.

Finally, in the special case of species symmetry (i.e., *λ*_1_ = *λ*_2_ = : *λ*, *α*_11_ = *α*_22_ = : *α*, *α*_12_ = *α*_21_ = : *β* and *b*_1_ = *b*_2_ = : *b*), *y* = 1 is solution of (S2.4) i.e., equal frequencies of both species. In this case, there are two other equilibria if and only if the derivative of (S2.4) is negative at *y* = 1. This derivative is given by

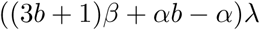
and is negative if and only if

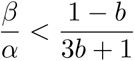

Equivalently, as *ρ* = *β/α* in this symmetric case, coexistence occurs if and only if

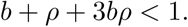

This is a special case of the conditions (S2.1) when *λ* ≫ 1

### General model: Carrying Simplices and Coexistence Criteria

To analyze the full model, we use the theory of monotone maps [Smith, 1998, Hirsch and Smith, 2005]. Define

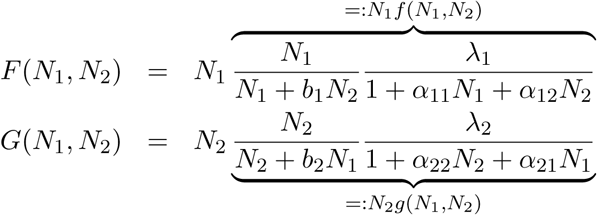

Then the dynamics of (2) is given by

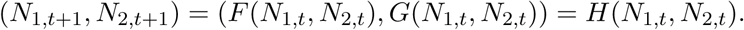

We recall a few definitions. The competitive ordering of the non-negative cone *C* = [0, ∞)^2^ is given by (*N*_1_,*N*_2_) ≥ *_K_* (*M*_1_,*M*_2_) if *N*_1_ ≥ *N*_2_ and *M*_1_ ≤ *M*_2_, (*N*_1_,*N*_2_) > *_K_* (*M*_1_,*M*_2_) if (*N*_1_,*N*_2_) ≥ *_K_* (*M*_1_,*M*_2_) and either *N*_1_ > *N*_2_ or *M*_1_ < *M*_2_, and (*N*_1_,*N*_2_) ≫ *_K_* (*M*_1_,*M*_2_) if *N*_1_ > *N*_2_ and *M*_1_ < *M*_2_. We will show that (2) is *strongly competitive*: *H*(*N*_1_,*N*_2_) ≫ *_K_ H*(*M*_1_,*M*_2_) whenever (*N*_1_,*N*_2_) > *_K_* (*M*_1_,*M*_2_). By Smith [1998, Proposition 2.1], *H* being strongly competitive follows from the derivative map 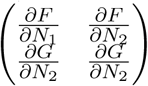 having the sign pattern 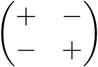. Taking the partial derivatives yields:

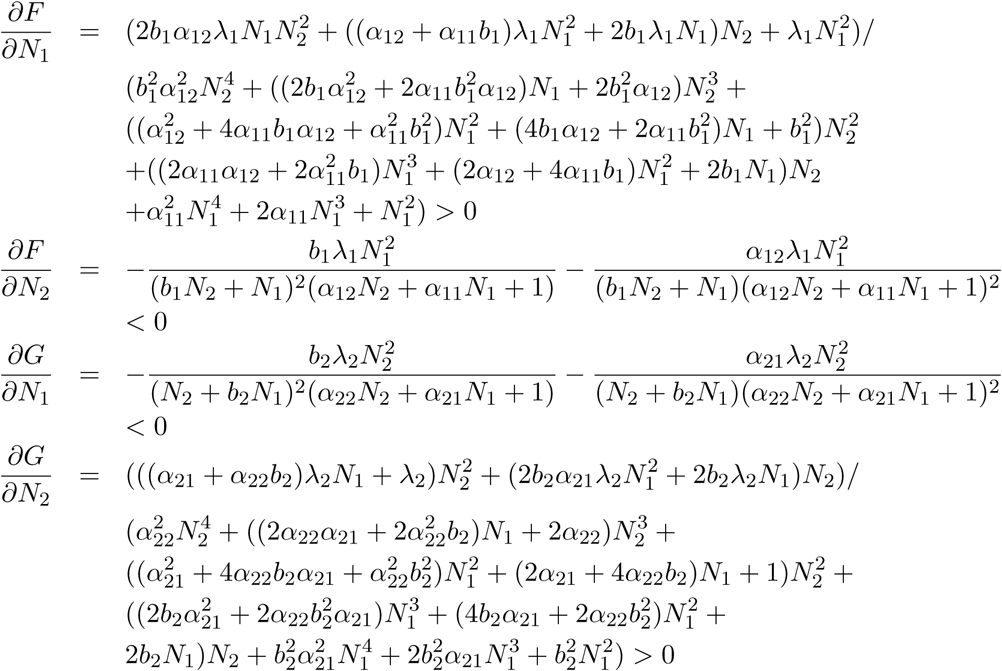

Hence, *H* is strongly competitive.

The sign of the determinant of *DH* is positive due to the following truly horrific calculation:

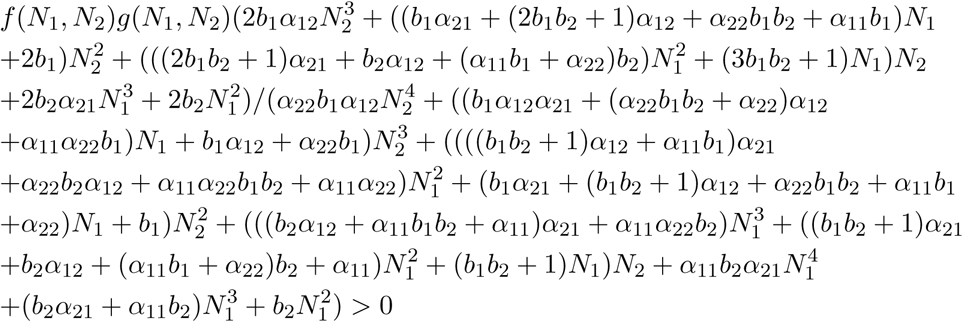

From these two calculations, we get that *DH* is invertible and all of its entries are strictly positive.

Using these facts, the work of [Smith, 1998] implies two key results. First, all solutions with *N*_1,0_*N*_2,0_ > 0 converge to a continuous (in fact Lipschitz), invariant curve Γ. This curve is known as the carrying simplex of the system. It has the important property that all radial lines in *C* intersect Γ in exactly one point. Thus, all possible frequencies of the species have a unique representation on this curve. One can view the dynamics of *H* restricted to Γ as describing the asymptotic frequency dynamics of the competing species. Second, all solutions of (2) converge to an equilibrium. We note: in the limiting case of *λ_i_* → ∞ for *i* = 1, 2, the simplified model (S2.3) can be viewed as a radial projection of the dynamics on the carrying simplex onto the standard simplex {(*N*_1_,*N*_2_): *N*_1_ ≥ 0,*N*_2_ ≥ 0,*N*_1_ + *N*_2_ = 1}.

As the boundary equilibria on Γ are stable, coexistence is possible if and only if there exist equilibria 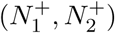 and 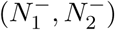 on Γ such that (i) 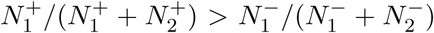 and (ii) there are no other equilibria 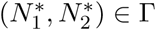 satisfying 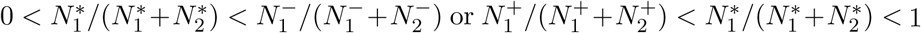 1. These equilibria correspond to the critical frequencies on Γ such that when the species lie on Γ between these critical frequencies, the species converge to an equilibrium supporting both species. Conversely, when the species initially lie on Γ outside these critical frequencies, the species dynamics converge to one of the boundary equilibria.

## Appendix S3 Stochastic Analysis

### Analysis of the simplified model (S2.2)

In the relative density coordinate system 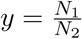, the stochastic simplified model takes the form

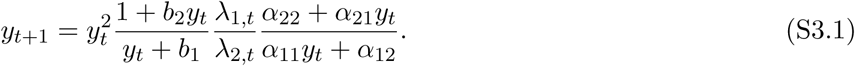

Assume that *b_i_* > 0, *α_i_* > 0, *β_i_* > 0, and (log *λ*_1,_*_t_*, log *λ*_2,_*_t_*) are a sequence of independent, identically distributed random vectors with a multivariate normal distribution with mean vector (*µ*_1_,*µ*_2_) and covariance matrix 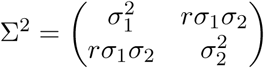 where *σ*_1_ > 0 and *σ*_2_ > 0.

If 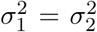 and *r* = 1, then *λ*_1,_*_t_/λ*_2,_*_t_* = exp(*µ*_1_ − *µ*_2_) is constant for all time *t*. Hence, the dynamics are deterministic and the analysis from Appendix S2 applies.

If 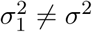 or *r<* 1, then log *λ*_1,_*_t_/λ*_2,_*_t_* is normally distributed with mean *µ*_1_ −*µ*_2_ and variance 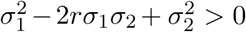. As *y* = 0 is a stable equilibrium, the proof of Roth and Schreiber [2014, Theorem 4.1] implies that for all *ε* > 0 there exists *δ* > 0 such that

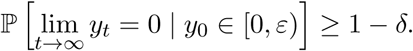

Namely, for initial values *y*_0_ sufficiently close to zero, the probability of losing species 1 is arbitrarily close to 1. Making the change of variables *z_t_* = 1*/y_t_*, the dynamics of (S3.1) become

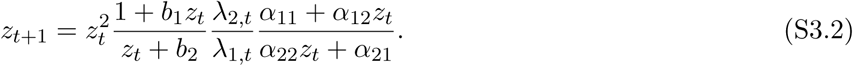

Reapplying the proof of Roth and Schreiber [2014, Theorem 4.1] implies that for all *ε* > 0 there exists *δ* > 0 such that

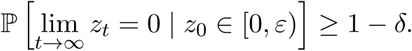

Namely, for initial values *z*_0_ sufficiently close to zero, the probability of losing species 2 is arbitrarily close to 1.

Since log *λ*_1,_*_t_/λ*_2,_*_t_* is normally distributed with positive variance, for any *M* > 0 there exists *γ* > 0 such that

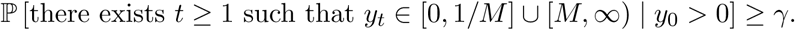

Consequently, the proof of Roth and Schreiber [2014, Theorem 3.2] implies that

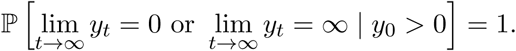

Hence, for any initial condition *y*_0_, with probability one, species 1 or species 2 goes extinct asymptotically. Moreover, whenever *y*_0_ > 0, there is a positive probability that either species goes extinct asymptotically.

### Verifying the storage effect for the model with survival

Consider the model with overlapping generations (i.e., *s_i_* > 0 for *i* = 1, 2) and no positive frequency-dependence (i.e., *b_i_* = 0 for *i* = 1, 2). As summarized in Ellner et al. [2016], the storage effect theory [Chesson, 1994] consider the instantaneous per-capita growth rate *r_i_* of species *i* written as a function environmental dependent parameter 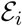 and the competition pressure 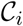. For our model, these dependencies are given as follows:

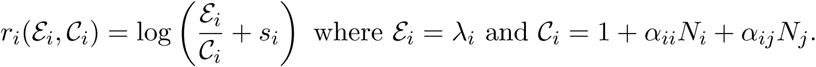

To get the storage effect, three ingredients are necessary [Ellner et al., 2016, Chesson, 1994]. First, the model needs to exhibit population buffering. This is satisfied if 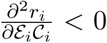 i.e., the effect of competition is weaker in years in bad years than in good years. Carrying out this second derivative yields:

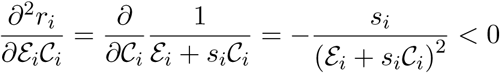
as desired. The second condition concerns positive stationary distributions 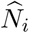 of the single species models *N_i_*(*t*+1) = *N_i_*(*t*)*λ_i_*(*t*)*/*(1+*α_ii_N_i_*(*t*))+*s_i_N_i_*(*t*). Benaïm and Schreiber [2009] implies such stationary distributions exist and are unqiue whenever 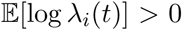 i.e., the species in isolation can increase when rare. Let 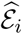 be the distribution of *λ_i_*(*t*) and 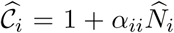. The storage effect requires that the covariance 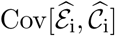 to be positive i.e., years which are good for species *i* also are years where they experience higher competition. When *λ_i_*(*t* − 1) and *λ_i_*(*t*) are uncorrelated, this covariance equals zero i.e., the current density of the species doesn’t influence the current environmental state. However, whenever *λ_i_*(*t* − 1) and *λ_i_*(*t*) are positively correlated, then *N_i_*(*t*) and *λ_i_*(*t*) tend to be positively correlated i.e., years where *λ_i_*(*t* − 1) is high tends to lead to high values of *N_i_*(*t*) and (due to the positive correlation) high values of *λ_i_*(*t*). Hence, 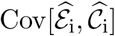 tends to be positive in this case. Finally, the storage effect requires that 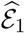 and 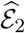 are not perfectly correlated and are sufficiently variable (i.e., each species has years in which they experience the better environment). This last condition is satisfied whenever the cross-correlation *r* in our model is less than one and the variance of log *λ_i_*(*t*) is positive for both species.

### Effects of temporal-autocorrelation on quasi-extinction risk

Figure S3-1 shows that increasing temporal autocorrelation increases quasi-extinction risk. Intuitively, with greater temporal autocorrelation, longer runs favoring one of the species is more likely. These longer runs result in the unfavored species decreasing in frequency and, consequently, more likely stay below a critical frequency at which their per-capita growth rate is less than one.

**Supplementary Figure S3-1:**
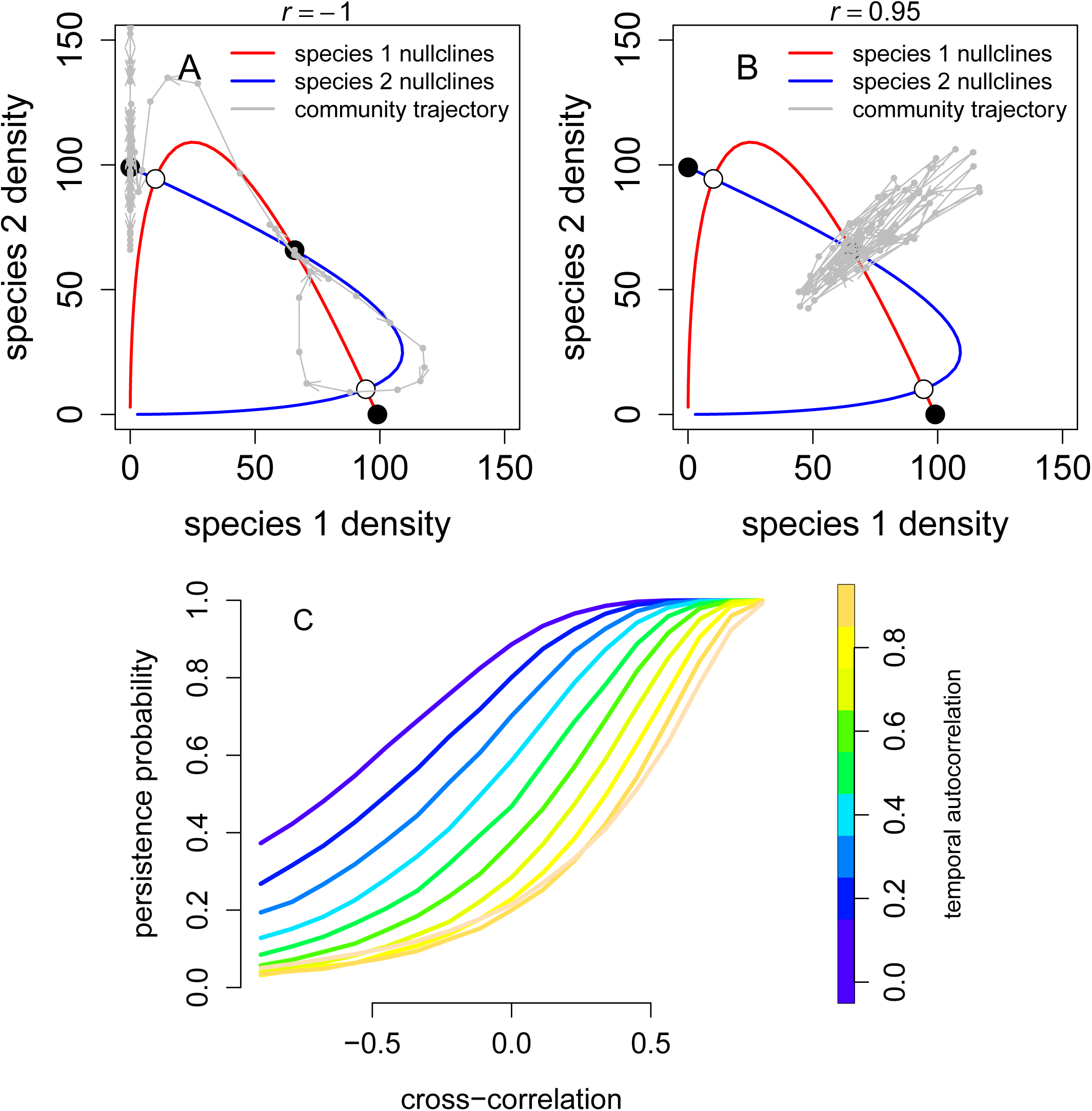
Positive temporal autocorrelation and negative cross-correlation increases quasi-extinction risk. In (A) and (B), simulations of the base model without survival are plotted in the phase plane with temporal autocorrelation *τ* = 0.5 and cross-correlations *r* as indicated. In (C), the probabilities of quasi-extinction in 50 years are shown for different values of the cross-correlation *r* and temporal autocorrelation *τ* without overlapping generations. Parameter values: *α*_11_ = *α*_22_ = 1, *α*_21_ = *α*_22_ = 0.2, *λ*_1_ = *λ*_2_ = 100, *b*_1_ = *b*_2_ = 0.1, *s*_1_ = *s*_2_ = 0, *σ*_1_ = *σ*_2_ = 0.5.

